# Global ecomorphological restructuring of dominant marine reptiles prior to the K/Pg mass extinction

**DOI:** 10.1101/2021.12.30.474572

**Authors:** Jamie A. MacLaren, Rebecca F. Bennion, Nathalie Bardet, Valentin Fischer

## Abstract

Mosasaurid squamates were the dominant amniote predators in marine ecosystems during most of the Late Cretaceous. Evidence from multiple sites worldwide of a global mosasaurid community restructuring across the Campanian–Maastrichtian transition may have wide-ranging implications for the evolution of diversity of these top oceanic predators. In this study, we use a suite of biomechanical traits and functionally descriptive ratios to investigate how the morphofunctional disparity of mosasaurids evolved through time and space prior to the Cretaceous-Palaeogene (K/Pg) mass extinction. Our results suggest that the worldwide taxonomic turnover in mosasaurid community composition from Campanian to Maastrichtian is reflected by a notable increase in morphofunctional disparity on a global scale, but especially driven the North American record. Ecomorphospace occupation becomes more polarised during the late Maastrichtian, as the morphofunctional disparity of mosasaurids plateaus in the Southern Hemisphere and decreases in the Northern Hemisphere. We show that these changes are not associated with strong modifications in mosasaurid size, but rather with the functional capacities of their skulls, and that mosasaurid morphofunctional disparity was in decline in several provincial communities before the K-Pg mass extinction. Our study highlights region-specific patterns of disparity evolution, and the importance of assessing vertebrate extinctions both globally and regionally. Ecomorphological differentiation in mosasaurid communities, coupled with declines in other formerly abundant marine reptile groups, indicates widespread restructuring of higher trophic levels in marine food webs was well underway when the K-Pg mass extinction took place.

## INTRODUCTION

Marine ecosystems were dominated by reptiles during the entire Mesozoic (Motani 2002; Massare 1987; Bardet 2012; Scheyer et al. 2014). Despite important turnovers at its base (Fischer et al. 2017; 2020), the Late Cretaceous is no exception, as mosasaurid squamates rapidly diversified (Bardet et al. 2008; 2007; Polcyn et al. 2014), achieving a cosmopolitan distribution prior to the Campanian (c.83.5 Mya) (Bardet et al. 2014; Polcyn et al. 2014), and colonised several ecological guilds until their global extinction at the K/Pg boundary mass extinction (66 Mya) (Martin et al. 2017). Prior to the Campanian, mosasaurid taxonomic richness saw a steep increase (Polcyn et al. 2014), with speciation in the Western Interior Seaway (WIS) in central North America triggering a diversification during the so-called ‘Niobraran Age’ (e.g. Kiernan 2002; Russell 1967). High taxonomic richness persisted through the mid-Campanian (e.g. Driscoll et al. 2019), where an abrupt taxonomic turnover is observed in central North American at the onset of the ‘Navesinkan Age’ (e.g. Russell 1993; Kiernan 2002). The abrupt shift observed in WIS mosasaurid community is mirrored in northern Europe (Lindgren 2004), Japan (Tanimoto 2005; Sato et al. 2012), South America (Jiménez-Huidobro, Simões, and Caldwell 2017), and to some extent in Oceania (Jiménez-Huidobro, Simões, and Caldwell 2017). Mosasaurids seem to maintain a high diversity throughout the Maastrichtian, yet with varying assemblages (Cappetta et al. 2014). Despite abundant remains, it is unknown whether these changes in taxonomic composition resulted in constriction of functional or ecomorphological variation of these top oceanic predators on provincial or global scales leading up to the end-Cretaceous mass extinction.

For the first time, we explore global mosasaurid ecomorphological variation throughout the final chapter of the Mesozoic (84–66 Mya) at both local and global scales, using a set of cranial and postcranial measurements, including data from several tens of high-precision 3D models. Because of the strong conservative forces governing mosasaurid bauplan evolution (e.g. hydrodynamic performance and phyletic heritage; e.g. Stubbs and Benton 2016), we did not anticipate significant temporal changes in craniodental morphofunctional disparity. Yet, we demonstrate polarisation of ecomorphospace occupation and significant drops in mosasaur disparity just before the K/Pg mass extinction, most notably in the Northern Hemisphere.

## MATERIAL AND METHODS

### Taxonomic and Morphological Sampling

Skull and jaw material from 93 mosasaurid specimens were collected, representing 56 species and all subfamilies and tribes (Polcyn et al. 2014; Simões et al. 2017). The taxonomic composition of mosasaurid clades in this study follows the results of Simões et al. (2017); we consider halisaurines as basal mosasaurids, and treat Russellosaurina (including Tethysaurinae, Tylosaurinae, Plioplatecarpinae, and Yaguarasaurinae) and Mosasaurina (including Mosasaurinae) as monophyletic groups. Morphometric information was collected from two main sources: three-dimensional laser and structured light surface scans as well as photogrammetric models were the preferred methodology, supplemented with two-dimensional published images and first-hand photographs. Laser scanned specimens were digitised using a Creaform HandySCAN 300 handheld laser scanner at resolution 0.2–0.5mm; structured light scanning was performed using an Artec Eva handheld scanner, at resolution 0.5mm; photogrammed models were captured using a Nikon D3000 DSLR camera (burst mode with light-ring), with 3D models generated using Agisoft Metashape 1.6.3., scaled in MeshLab 2020.06 (Cignoni et al. 2008). All specimens are listed in Supplementary Information 1: Specimen List; 3D models will be provided in the open-source repository MorphoSource on peer-reviewed acceptance.

### Linear Measurements and Functional Ratios

Twenty-four linear measurements were taken across the entire skull including dentition (Figure 1). Linear measurements on 3D scans were taken using the measurement tool in MeshLab 2020.06; measurements on 2D images were performed using the line measuring tool in ImageJ (Schneider, Rasband, and Eliceiri 2012). These measurements were then used to generate 16 functional ratios describing the craniodental architecture (Table 1). All these traits have clearly established functional importance or outcomes (Figure 1; for further details, see Supplementary Information 2: Functional Ratios). Examples include the ratio of mandibular lever arms (proxies for mechanical advantage, i.e. ratio of muscular input force to output force on prey items), supratemporal fenestrae area (proxy for cross-sectional area of combined jaw adductor musculature), and relative orbit size (amount of the skull dedicated to housing the eyeball).

**Table 1.**
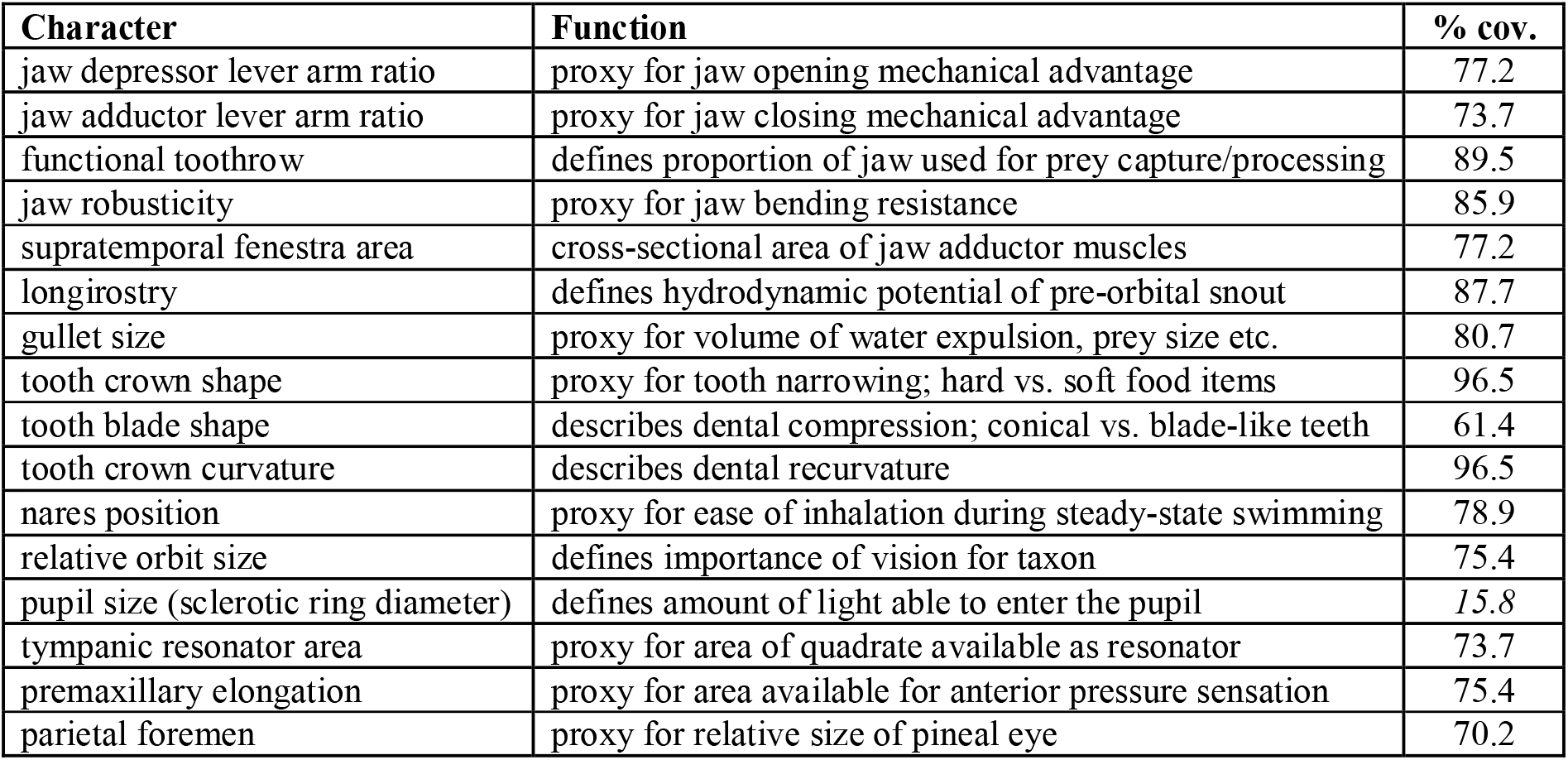
Functional traits derived from linear measurements of mosasaurid skulls and jaws. Definitions, calculations and diagrammatic representations for each trait can be found in the Supplementary Information 3: Functional Ratios. % cov. = percentage of specimens represented by each trait. Percentages in italics fall outside the completeness threshold of 40%.

**Figure 1.**
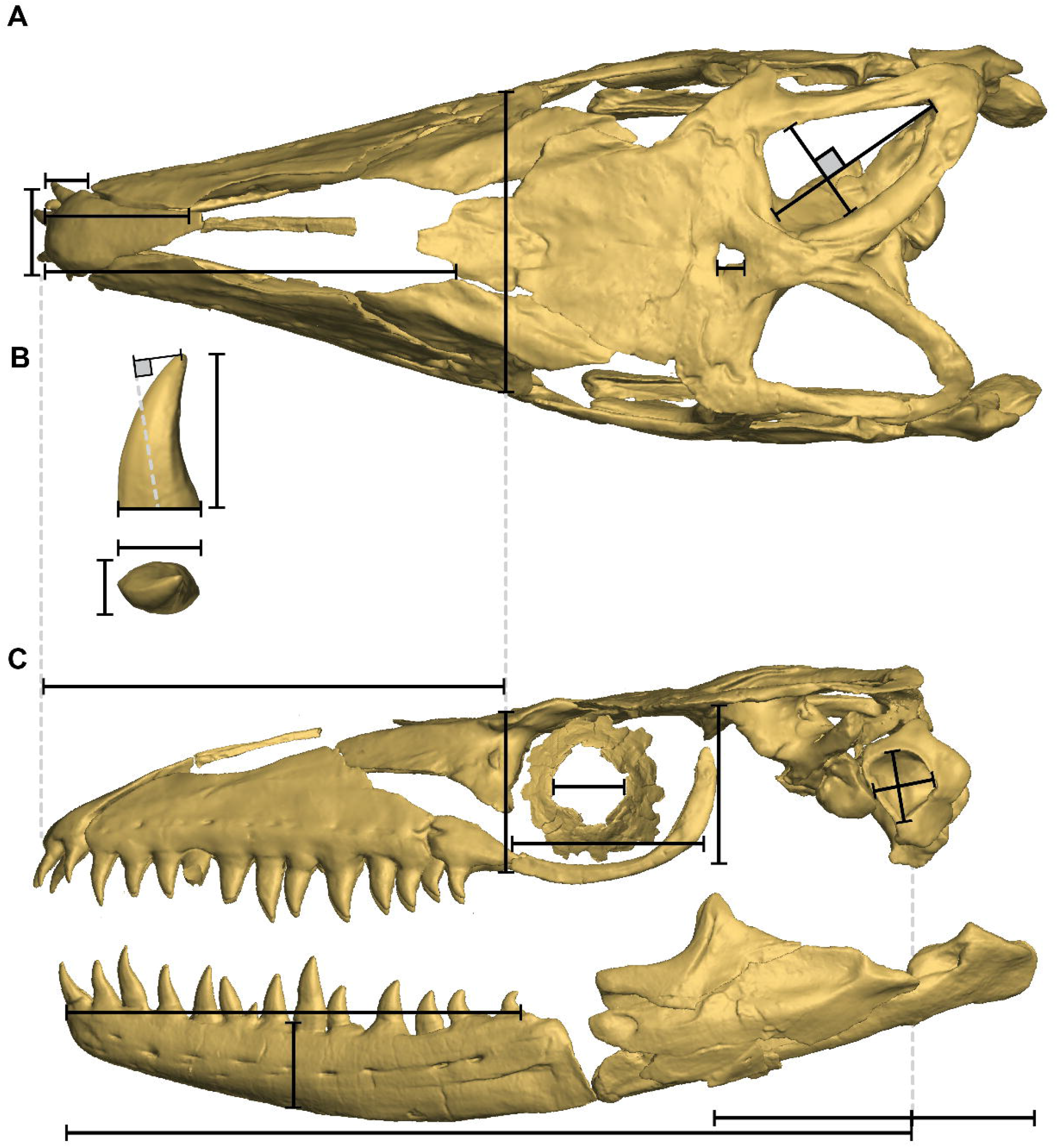
Linear measurements of the mosasaurid skull used for quantitative trait comparisons and disparity analyses. Measurements on the skull and exemplar dentition are shown: **(A)** skull in dorsal aspect, **(B)** dentition in lateral and occlusal aspect, (C) skull in left lateral aspect. Filled squares denote measurements taken perpendicular to one another or to the edge of a bone. Black lines describe measurements used for trait quantification; dotted grey lines are used to clarify where specific measurements are recorded from and to. Functional traits and their ecomorphological importance are presented in Table 1. Models based on IRSNB R33b *Prognathodon solvayi*.

A threshold of 40% trait completeness was applied to the sample, on each specimen; percentages of missing data per species can be found in Supplementary Information 3: Species Coverage. A resultant craniodental dataset with all 16 functional ratios was subjected to the 40% threshold. Following the threshold computations, 18.2% of missing trait data was recorded, and the dataset included all 58 taxa studied. Trait ratios were standardised using a z-transformation to assign all characters a mean of 0 and a variance of 1; data which cleared the 40% completeness threshold were used to compute a Euclidean distance matrix for ordination analyses and disparity calculation.

### Ecomorphospaces

Ordination of trait data was visualised in two-dimensions in two ways: a principal coordinates analysis (PCoA) and a non-metric multidimensional scaling approach (NMDS). PCoAs were performed using a cailliez correction criterion to correct for negative eigenvalues (using ape v.5.3; Paradis, Claude, and Strimmer 2004), and were preferred to principal components analysis (PCA) as PCoA allows missing values in the Euclidean distance matrix. Comparisons between PCoA and non-metric multidimensional scaling (NMDS) ordination demonstrated comparable patterns of ecomorphospace occupation. NMDS are used for visualisation here as they pack more variation of the data into a two-dimensional graph. However, as NMDS axes are not ideal to use as variables for disparity analyses because of their non-metric properties, all PCoA axes were chosen for assessment of morphofunctional disparity in through time, within clades, and across geographic regions. Comparative NMDS analyses were performed in ‘vegan’ v.2.5-6 (Oksanen et al. 2018); graphical results from PCoA ecomorphospaces can be found in Supplementary Figure S1. Kernel 2D density estimates were used to visualise density-based macroevolutionary landscapes, plotted onto NMDS ecomorphospaces, following the methodology of Fischer et al. (2020). Moreover, mandible length (proxy for body size) was used both in scaling datapoints to visually inspect the spread of large-sized mosasaurids, and additionally to compare the spread of body sizes in mosasaurids through the Campanian-Maastrichtian.

### Disparity

Morphofunctional disparity was calculated based on PCo axes. In addition, mosasaurid communities from four geographically distinct regions were investigated: the Western Interior Seaway (WIS); Northern Tethys Province (NTP); Southern Tethys Province (STP; Bardet 2012); and Weddellian Province (consisting of South-East Oceania, the Antarctic peninsula and Patagonia; WP). Disparity was measured through time, focusing on time bins bearing mosasaurid fossils during the latest Cretaceous: Early Campanian (83.60–77.85 Mya); Late Campanian (77.85–72.10 Mya); Early Maastrichtian (72.10–69.05 Mya); Late Maastrichtian (69.05–65.50 Mya). The focus was made on these time periods as they encompassed the mosasaurid taxonomic turnover in the mid-Campanian, and enabled the investigation of disparity in the lead up to the end-Cretaceous mass extinction. Total disparity within mosasaurid clades and geographic regions during these time periods was also calculated. Disparity analyses were performed in RStudio using the dispRity package (v.1.5.0) (Guillerme 2018). The sum of variances (SoV) disparity metric was preferred, as it demonstrates robusticity to sample size variation between time bins (Ciampaglio, Kemp, and McShea 2009). Alternative disparity metrics (pairwise dissimilarity, PD; sum of ranges, SoR) were also tested to corroborate patterns observed using SoV metric (Supplementary Table S1). Bootstrap iterations were set at 1000 repetitions; additional bootstrapping procedures were performed to account for false positive results when testing for significant differences (see Supplementary Table S1). Here, we adapt the terminology from population ecology to assess disparity at the regional level (here termed ‘α-disparity’) and global level (‘γ-disparity’). In order to examine how mosasaurid ecomorphological disparity was differentiated across regional communities, γ-disparity per time bin was divided by mean α-disparity across all provinces per time bin (following population ecology methods e.g. Legendre 2008), creating ‘ß-disparity’. Beta-disparity can be defined as a measure of disparity differentiation; a high ß-disparity indicates a greater range of mean α-disparity values within a specific time-bin, whereas low ß-disparity indicates more uniformity in mean α-disparity values, suggesting less ecomorphological differentiation between disparities across communities. Mean bootstrap estimates (1000 replications) were used as metrics for α- and γ-disparity calculations, and compared through four time-bins (Early Campanian; Late Campanian; Early Maastrichtian; Late Maastrichtian). Changes in disparity between subsequent time bins and between clades/geographic regions were tested for using non-parametric Wilcoxon tests in the ‘stats’ package (v.4.0.3) (R Core Development Team 2008), with Bonferroni corrections for multiple comparisons.

## RESULTS

### Morphospace occupation

We quantified mosasaurid craniodental disparity across the Campanian-Maastrichtian interval. Our aims were to establish whether faunal transitions yielded changes in disparity in mosasaurids as well as whether global and provincial mosasaurid disparity was in decline prior to the end-Cretaceous mass extinction. Most notably, large ‘megapredatory’ taxa with cutting dentition, including almost all tylosaurines (Figure 2A; filled purple squares), and the majority of large mosasaurines cluster in the ecomorphospace (Figure 2A). These results suggest that overall skull functional morphology within these two (occasionally contemporaneous) clades of megapredatory marine reptiles converged, despite numerous phylogenetic differences. A number of brevirostrine mosasaurines occupy regions of positive NMDS axis 1, typified by relatively large supratemporal fenestrae, deep jaws, blunt rostra and crushing dentition (e.g. *Globidens* spp.). The upper half of ecomorphospace (positive values along NMDS axis 2) is occupied predominantly by primitive mosasaurids (halisaurines, tethysaurines) and *Plioplatecarpus* spp., which all had large orbits, gracile skulls, and recurved, piercing teeth (Figure 2A; see also Supplementary Figure S1); they constitute our ‘grasping’ cluster (Figure 2A & 2E).

**Figure 2.**
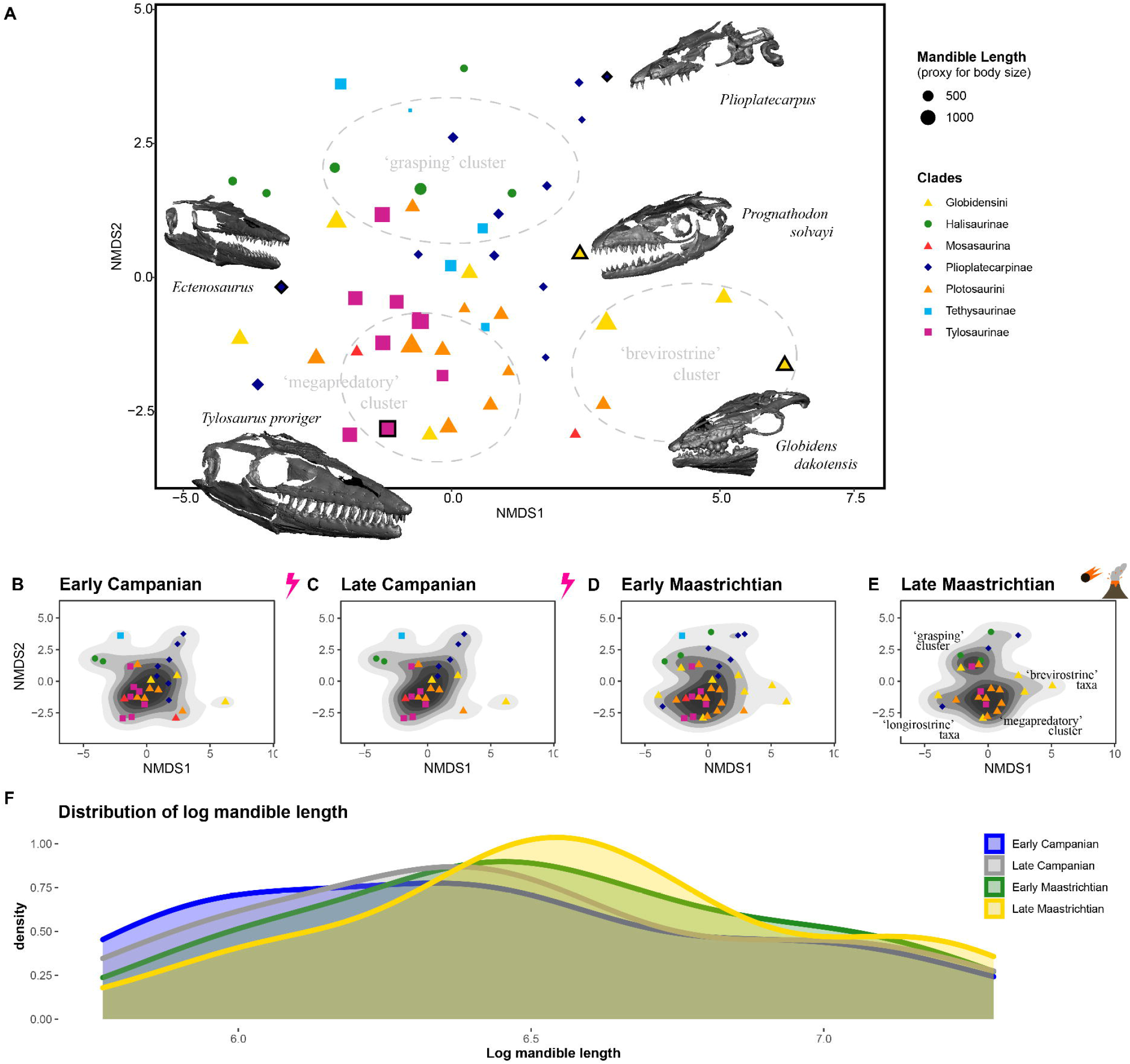
Functional ecomorphospace and size distribution in mosasaurids. Functional ecomorphospace occupation (based on NMDS axes) by all mosasaurids in the sample (**A**) with ecomorphological clusters and representative 3D models of skulls. Data points outlined in bold represent the placement of exemplar skulls; data point size represents relative skull size (based on mandible length). Functional ecomorphospace for each time bin through the Campanian-Maastrichtian (**B-E**) demonstrating changes in density and isolation of ecomorphological clusters in Late Maastrichtian (**E**). Size distribution of mosasaurids through the Campanian-Maastrichtian (**F**) demonstrating shifts in the density of small and mid-sized mosasaurids from Early Campanian (ECam) to Late Maastricthian (LMaa).

The density of ecomorphospace occupation though time (Figure 2B-E) reveals a series of changes across the Campanian-Maastrichtian interval. Many mosasaurines and russellosaurines occupy a large ‘megapredatory’ region through the Early Campanian to the Early Maastrichtian. The majority of russellosaurines disappear afterwards, strongly altering the pattern of ecomorphospace occupation (Figure 2E) by creating a clear divide between two main clusters in the late Maastrichtian bin: the ‘megapredatory’ cluster, formed predominantly by *Tylosaurus* and *Mosasaurus* and the ‘grasping’ cluster, formed by *Plioplatecarpus*, halisaurines, and the Weddellian taxa *Taniwhasaurus oweni* and *Rikisaurus tehoensis*. In addition to this polarisation of mosasaurid craniodental shape, a few taxa also evolved longirostrine (e.g. *Gavialimimus*) and brevirostrine (likely durophagous; e.g. *Globidens* spp.) morphologies (Figures 2A-B). Our analyses of clade disparity add support to the ecomorphospace signal, with decreases in tylosaurine, plioplatecarpine, and plotosaurin disparity through the Maastrichtian (Supplementary Figure S2).

### Evolution of skull size

Changes in mosasaurid communities also resulted in slight variation in skull size distributions (proxy for body size) across the Campanian – Maastrichtian interval (Figure 2F). Early Campanian size distribution is notably more uniform, with comparable densities of large and small mosasaurids (Figure 2F; blue line). By comparison, density of smaller species is lower in the Late Maastrichtian, leading to a peak in mid-sized and very large species (Figure 2F; yellow line). This pattern tracks the presence of multiple very large late Maastrichtian tylosaurines and plotosaurins (a pattern mirrored in sharks; Cappetta et al. 2014), and highlights the extinction of smaller species which were abundant during the Campanian (e.g. *Clidastes, Plesioplatecarpus*) (Figure 2B-C). However, these differences are not significant, indicating that the changes in ecomorphospace occupation and disparity we observe are not associated with strong changes in mosasaurid size, but rather with the functional capacities of their skulls.

### Spatiotemporal evolution of disparity

We find a significant increase in global (γ) ecomorphological disparity in mosasaurids (Figure 3A) coincident with taxonomic turnovers known to have occurred at the mid-Campanian boundary (the ‘Niobraran-Navesinkan’ transition; Lindgren 2004; Tanimoto 2005; Sato et al. 2012; Jiménez-Huidobro, Simões, and Caldwell 2017; Kiernan 2002; Russell 1993). The observed increase in γ-disparity is common across all disparity metrics we computed (sum of variances, sum of ranges, pairwise dissimilarity; Supplementary Table S3). Our results demonstrate that γ-disparity of mosasaurid ecomorphologies increased from Early to Late Campanian and continued increasing until the Early Maastrichtian, mirroring the expansion of the craniodental ecomorphospace occupation (Figure 2B-D). Mosasaurid diversity (in this sample) somewhat tracks fluctuations in disparity, but not to the same magnitude (Figure 3). By the Late Maastrichtian, γ-disparity is higher than that recorded throughout the Campanian, despite fewer species being present in the Late Maastrichtian (Figure 3A; Table 2).

**Table 2.** Mosasaurid population disparity (alpha, gamma and beta) from Early Campanian to Late Maastrichtian. Sum of variances metric used. Highest mean disparity values highlighted in bold. WIS = Western Interior Seaway; NTP = Northern Tethys Province; STP = Southern Tethys Province; WP = Weddellian Province.

| Disparity |  | Early Campanian | Late Campanian | Early Maastrichtian | Late Maastrichtian |
| --- | --- | --- | --- | --- | --- |
| gamma ($\gamma$) | | 18.90 | 19.30 | **21.19** | 20.12 |
| alpha ($\alpha$) | Mean | 15.94 | 16.17 | **19.26** | 14.63 |
|  | WIS | *18.43* | **19.57** | *18.52* | 9.76 |
|  | NTP | *22.03* | *21.60* | **23.35** | *17.50* |
|  | STP | *10.22* | *10.44* | **22.04** | *21.51* |
|  | WP | *13.07* | *13.06* | **13.13** | 9.75 |
| beta ($\beta$) | | 1.186 | 1.194 | 1.100 | **1.375** |

**Figure 3.**
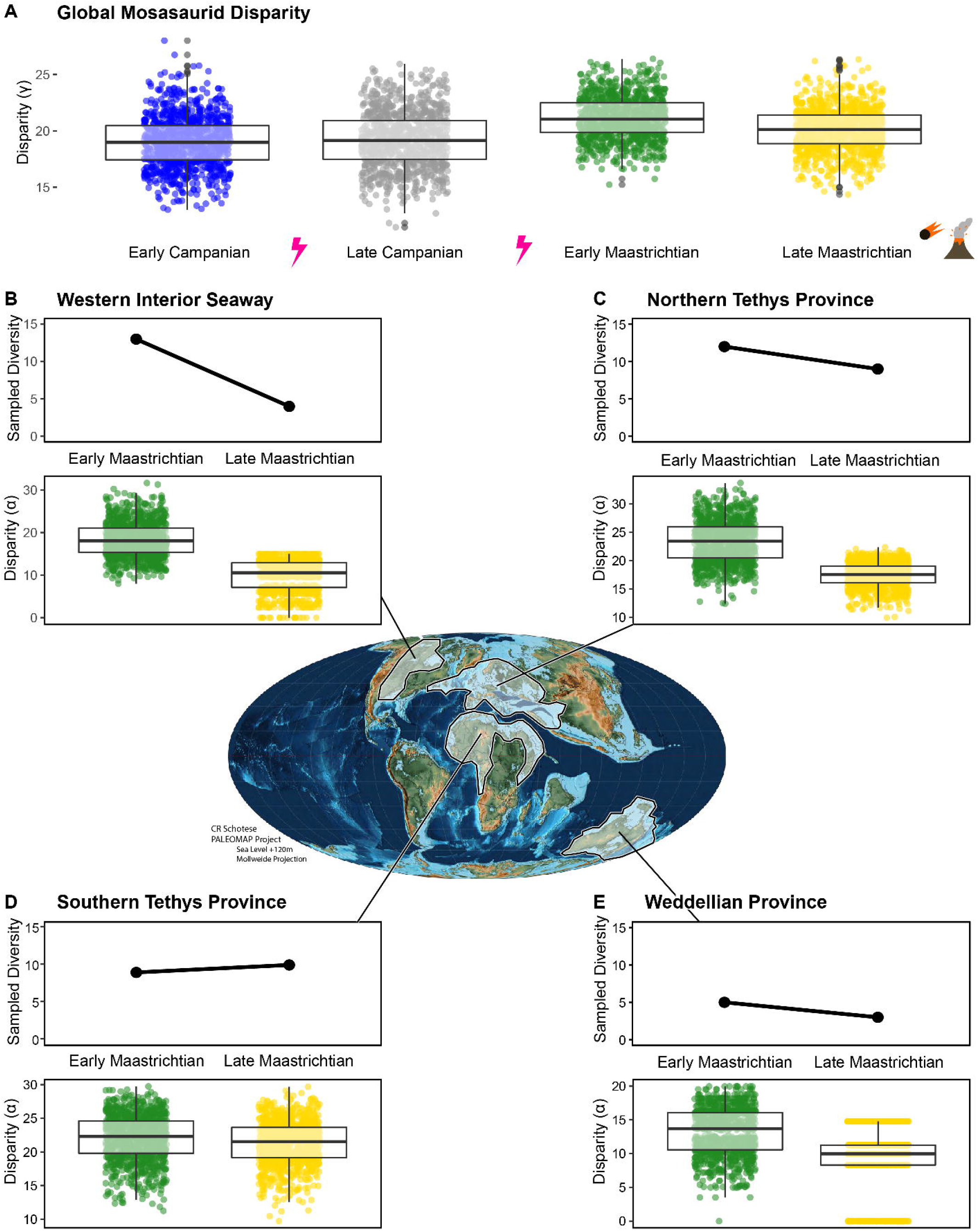
Global and provincial mosasaurid craniodental disparity through time. Global (γ) craniodental disparity of mosasaurids from Early Campanian to Late Maastrichtian (**A**) presented alongside raw sample diversity and provincial (α) disparity for the Maastrichtian of the Western Interior Seaway (**B**), Northern (**C**) and Southern (**D**) Tethys Provinces, and Weddellian Province (**E**). Approximate extent of provincial regions projected onto palaeomap, estimated for mid-Maastrichtian (72 Ma). Disparity estimates were generated using the sum of variances (SoV) metric; significant differences were recovered between all sequential time bins for both global and provincial datasets using pairwise Wilcoxon testing (see also Supplementary Table S1 & S3). Palaeomap provided by CR Schotese (PALEOMAP atlas for ArcGIS) (Scotese 2014).

While disparity increases on global (γ) and provincial (α) scales from the Campanian to the Maastrichtian (Table 2), we observe significant declines in γ-disparity from early to late-Maastrichtian leading up to the K-Pg mass extinction (Figure 3A-E). When examined at the provincial level (Figure 3B-E), the early–late Maastrichtian transition records declines in α-disparity for all provinces (with the exception of STP; Table 2) using almost all disparity metrics, reinforcing the global outlook of a significant decline in ecomorphological disparity in mosasaurids in the latest Maastrichtian. This disparity decrease is found within all well-sampled clades as well, no matter the disparity metric used, with the exception the pairwise dissimilarity metric which slightly increased for the hyper-disparate group Globidensini through the Maastrichtian (Figure 2D-E). Sharp decreases in tylosaurine and plioplatecarpine presence in the Western Interior Seaway contribute to the decline in α-disparity in this region; by contrast, the presence of highly disparate globidensins in the Southern Tethys Province contributes toward more stable overall γ-disparity during the Maastrichtian (Figure 3A & 3D). Differentiation of disparity across regions (i.e. β-disparity) increases from early to late Maastrichtian (Table 2); this increase in β-disparity can be attributed to several factors: decreases in α-disparity in several (but crucially, not all) observed provinces; reductions in taxon count (Figure 3B-E); decreased occupancy of previously commonly exploited niches (e.g. reduction of ‘megapredators’; Figure 2E); and increased endemism (e.g. Moroccan fauna of the STP; Strong et al. 2020; Lingham-Soliar 2002; Leblanc, Mohr, and Caldwell 2019; Longrich, Bardet, Khaldoune, et al. 2021; Longrich, Bardet, Schulp, et al. 2021).

## DISCUSSION

### The necessity for regional and global assessments of pre-extinction diversity and disparity

The influence of localised faunal assemblages in the fossil record is well known to affect global patterns diversity and disparity (i.e. Close et al. 2019; Upchurch et al. 2011; Condamine et al. 2021; etc.). For many groups of large tetrapods, the global fossil record is not well resolved, whereas regional sampling in certain geographic areas is strong, and consequent global biodiversity/disparity estimates can be heavily reliant on those few regions (e.g. Brusatte et al. 2012; Sax and Gaines 2003; Benson et al. 2010; Cleary et al. 2015; etc.). In many studies, including ours, it is clear that the sampling effort in North America over the past 150 years is an important factor in estimating pre-Maastrichtian tetrapod diversity and disparity (e.g. Maidment et al. 2021; Vavrek and Larsson 2010; Longrich, Scriberas, and Wills 2016). Regional diversity patterns are thus likely to contain an important signal, as the highly-fragmented world of the Mesozoic and Cenozoic likely resulted in ecosystems with distinct compositions and physico-chemical parameters (Zaffos, Finnegan, and Peters 2017). This reality has often been overlooked when analysing tetrapod diversity and disparity patterns leading up to and across the K/Pg mass extinction. Indeed, a series of studies on the extinction of non-avian dinosaurs have recovered conflicting results (Brusatte et al. 2012; Condamine et al. 2021; Maidment et al. 2021; Dean, Chiarenza, and Maidment 2020; Chiarenza et al. 2019; Vavrek and Larsson 2010), notably because of their varying treatment of regional differences and their sampling.

Even though the fossil record of mosasaurids appears only weakly biased (Driscoll et al. 2019), and marine reptile sampling indicators are generally excellent for the Campanian-Maastrichtian interval (Fischer et al. 2016), our results clearly indicate regional variations in the ecomorphological disparity patterns of mosasaurids. The drivers of these differences should not necessarily be regarded as global; a telling example are the provincial disparity patterns during the Maastrichtian (Figure 3B-E), which may be associated with the magnitude of the environmental changes resulting from the sea level regressions. Indeed, the epicontinental WIS greatly changed in extent and shape during the Maastrichtian (Berry 2017; Slattery et al. 2013), and this region records the steepest decrease in α-disparity, while deeper basins such as Northern and Southern Tethys Provinces were seemingly less affected (Bardet et al. 2014; Hornung, Reich, and Frerichs 2018; Lindgren 2004; Bardet 2012). In this context, focussing on the abundant North American record to reconstruct the global diversity or disparity patterns of mosasaurids would result in a steeper late Maastrichtian decrease than that which was computed for other regions, hence confounding regional and global factors at play prior to the K/Pg mass extinction.

### Pre-K/Pg mosasaurid turnovers and crises

The ‘Niobraran-Navesinkan’ (Early to Late Campanian) taxonomic transition from russellosaurine-to-mosasaurine-dominated communities was initially identified in the Western Interior Seaway (WIS) (e.g. Russell 1993), with similar turnovers identified in multiple other regions across the globe (Lindgren 2004; Tanimoto 2005; Sato et al. 2012; Jiménez-Huidobro, Simões, and Caldwell 2017). We show that, far from experiencing a global γ-disparity decline during this turnover, mosasaurids significantly increased in disparity across this transition. However, this is in no small part due to the extinction of more ‘generalist’ and small-sized plioplatecarpines and tylosaurines (Figure 3A). These extinctions in the WIS reduced the density of ‘megapredatory’ and ‘generalist’ ecomorphologies in the Late Campanian bin (Figure 2B-C), causing increased polarisation of the remaining ecomorphologies exhibited by mosasaurid taxa. The removal of common morphologies from a sample can impact disparity as much as the inclusion of highly disparate forms; this is the case for a series of disparity metrics (but obviously not the sum of ranges). We observe this pattern for mosasaurid γ-disparity through the Campanian (Figure 2B-C; Figure 3A). Actually, the increase in morphofunctional disparity at the ‘Niobraran-Navesinkan’ transition is a phenomenon local to the Western Interior Seaway, with a high enough amplitude to influence global patterns; other regions maintain stable morphofunctional disparity through this interval (Figure 3; Table 2). The cause behind the abrupt shift in mosasaurid community composition across the ‘Niobraran-Navesinkan’ is as yet unclear. A decrease in oceanic temperature between the mid- and late-Campanian is coincident with the turnover (Polcyn et al. 2014; Linnert et al. 2016), which essentially removed smaller species within multiple clades (e.g. *Selmasaurus, Plesioplatecarpus, Tylosaurus kansasensis*). This seemingly selective extinction suggests that body size was an important factor in lineage survival across this local event (Figure 2B-C and 2F). By contrast, mosasaurid assemblages from the Campanian elsewhere do not suggest clear reductions in smaller species at the mid-Campanian boundary (Lindgren 2004; Jagt 2005), nor do they exhibit a radical shift in α-disparity (Table 2).

Later, a decrease in global γ-disparity and α-disparity is found within the Maastrichtian, in nearly all regions and across all clades. When the differentiation of ecomorphological disparity between geographical regions is considered (i.e. β-disparity; Table 2), it is clear that the Late Maastrichtian was a time of increased regionalisation of mosasaurid disparity, rather than a consistent global decline. Communities of mosasaurids in the WIS, NTP and WP are shown to be notably more homogeneous in the Late Maastrichtian than those of the Early Maastrichtian (Figure 3B-E), with the WIS and WP communities represented by very few taxa within only two tribes (Plotosaurini+Globidensini, and Plotoaurini+Tylosaurini respectively). By contrast, the Late Maastrichtian mosasaurid community of the STP was comprised of a ecomorphologically diverse assemblage of globidensins (e.g. *Globidens*), plotosaurins (*Mosasaurus* spp.), halisaurines (e.g. *Pluridens*), and plioplatecarpines (e.g. *Gavialimimus*) (Strong et al. 2020; Leblanc, Caldwell, and Bardet 2012; Bardet et al. 2004; Longrich, Bardet, Schulp, et al. 2021), yielding a high α-disparity in this region (Figure 3D). The retention of disparate ecomorphologies of STP mosasaurids through the Maastrichtian drives the spike in β-disparity observed in the Late Maastrichtian (Table 2), representing a peak in provincial differentiation. The predominantly bimodal landscape of mosasaurids in the late Maastrichtian (Figure 2E) suggests that, while a variety of niches were still being occupied by low densities of disparate mosasaurids, numerous Northern and Southern Tethys mosasaurids exhibited ‘megapredatory’ or ‘grasping’ functional adaptations (Figure 2; also Bardet 2012; Lindgren 2004; Bardet et al. 2014). Becoming increasingly apparent is the importance of the Southern Tethys Province (including Afro-Arabia, Morocco, Niger-Nigeria, Angola and eastern Brazil) in estimating late-Maastrichtian marine reptile diversity and disparity (e.g. Strong et al. 2020; Leblanc, Mohr, and Caldwell 2019; Longrich, Bardet, Schulp, et al. 2021; Longrich, Bardet, Khaldoune, et al. 2021). Understanding patterns such as these are vital for the accurate interpretation of faunal dynamics and functional variation before and after extinction events. If only γ-disparity of mosasaurids were considered, then this group could be interpreted as being in decline prior to their ultimate demise at the K/Pg boundary. However, it is apparent that when both α- and β-disparities are taken into account, some regional communities were most certainly declining in taxonomic diversity and ecomorphological disparity, whereas others were only minimally affected on both counts.

### How selective are the pre-K/Pg extinctions in marine reptiles?

The Early Maastrichtian is identified here as the time of greatest γ-disparity of mosasaurids (Figure 3A), with expansion of ecomorphospace occupation by longirostrine and brevirostrine ecomorphologies, in addition to an increase in halisaurine and plioplatecarpine species occupying new regions of ecomorphospace in the ‘grasping’ cluster (Figure 2A+D). This Early Maastrichtian rise in disparity appears to be driven by multiple originations in the two Tethys Provinces (North and South). With regards to other marine reptile fauna, ichthyosaurians and pliosaurids were long gone by the Late Maastrichtian (Schumacher 2011; Fischer et al. 2016; Bardet 1994), and the predominant marine reptiles in the Campanian-Maastrichtian were restricted to xenopsarian plesiosaurians, mosasaurids, chelonioids, and crocodylians (e.g. Bardet et al. 2014). These clades do not seem to follow a disparity pattern similar to that recovered here for mosasaurids. For example, polycotylid plesiosaurians were already in decline in both phylogenetic diversity and ecomorphological disparity during the Campanian-Maastrichtian interval (Fischer et al. 2018), with only the polycotyline *Dolichorhynchops herschelensis* and the occultonectian *Sulcusuchus erraini* potentially being present during the Maastrichtian (in addition to a handful of indeterminate remains; Sato 2005; Kaddumi 2006; O’gorman and Gasparini 2013; Fischer et al. 2018). Robust evaluations of elasmosaurid disparity are still lacking, but the range of phenotypes (either in terms of phylogenetic diversity, osteology or relative neck length) appears to still be broad during the Maastrichtian, although within-Maastrichtian changes have not yet been computed (Fischer et al. 2021). Similarly, within-Maastrichtian disparity dynamics have not been explored for marine testudines, although previous assessments of testudines indicate a peak in cranial morphological variation in the Maastrichtian (Foth and Joyce 2016; Foth, Ascarrunz, and Joyce 2017), and a consistent contribution to the diversity of feeding morphologies across marine reptiles from Campanian to Maastrichtian (Stubbs and Benton 2016). By contrast, aquatic crocodyliformes exhibit comparatively low disparity in the latest Cretaceous. Dramatic declines in marine crocodyliform morphological disparity are known to have occurred from Jurassic to Cretaceous, which potentially left niches open for mosasaurids to exploit during their own radiation in the Cenomanian (Stubbs et al. 2013; Stubbs and Benton 2016). Subsequently, crocodyliform morphological disparity was in decline throughout the late Cretaceous (Wilberg 2017; Stubbs et al. 2021); the exception to this pattern were the aquatic dyrosaurid tethysuchians, which exhibited a rapid burst of morphological evolution in the Maastrichtian (Stubbs et al. 2021), coinciding with increased regionalisation of mosasaurid ecomorphological disparity revealed in this study. The increased endemism and expansion into ‘grasping’ and ‘longirostrine’ ecomorphologies by mosasaurids in the Late Maastrichtian combines with isotopic analyses of trophic structure in the Maastrichtian of the Southern Tethys Province (Martin et al. 2017), suggesting increased dietary specialisation, as multiple predators coexisted and often fed upon prey from a single trophic level. Patterns of taxonomic diversity and ecomorphological disparity across multiple marine reptile groups indicate that wholesale (and in some cases fragile) restructuring of marine trophic webs was underway before the K-Pg mass extinction event.

## Acknowledgements

The authors would like to thank the museum curators for access to their specimens for photographing and scanning: Megan Sims, Anna Whitaker, and Chris Beard (KUVP, Kansas); Chase Shelburne and Laura Wilson (FHSM, Kansas); Amanda Millhouse and Matt Miller (USNM, Washington DC); Amy Henrici (CMNH, Pittsburgh); Bill Simpson (FMNH, Chicago); Laura Vietti (UW, Laramie); Jodee Reed (FFHM, Oakley); Annelise Folie (RBINS, Brussels); Ronan Allain (MNHN, Paris); Benjamin Kear (PMU, Uppsala). We also thank Mike Polcyn for access to 3D data from published articles. This work was funded by grants from the Fonds De La Recherche Scientifique (F.R.S.-FNRS): MIS F.4511.19 “SEASCAPE” (VF), FNRS travel grant 35706165 (JAM), and a FRIA doctoral fellowship (RFB).

## Author Contributions

JAM and VF conceived the study. JAM, RFB, and VF collected raw data. JAM and VF conducted statistical analyses and designed figures. JAM, NB, and VF wrote the manuscript. All authors contributed to the final manuscript.

## Notes

### Competing Interest Statement

The authors have declared no competing interest.

