## Supplementary Information for "Global ecomorphological restructuring of dominant marine reptiles prior to the K/Pg mass extinction"

**Table of Contents:**

1. Specimen List
2. Functional Ratios
3. Species Coverage
4. Supplementary Figures
5. Supplementary Tables
1. **Specimen List**

| **Specimen No.** | **Genus** | **Species** | **Reference** |
| --- | --- | --- | --- |
| SGMA 12/60 | *Angolasaurus* | *bocagei* | Lingham-Soliar 1994 |
| IRSNB R 43 | *Carinodens* | *belgicus* | Bardet et al. 2014 |
| ANSP 10193 | *Clidastes* | *propython* | Cope 1871 |
| FHSM VP 17576 | *Clidastes* | *propython* | laser scan |
| FHSM VP 2071 | *Clidastes* | *propython* | photo |
| FMNH PR 495 | *Clidastes* | *propython* | structured-light scan |
| KUVP 1000 | *Clidastes* | *propython* | laser scan |
| KUVP 1022 | *Clidastes* | *propython* | laser scan |
| USNM 11719 | *Clidastes* | *propython* | laser scan |
| UW 15558 | *Clidastes* | *propython* | laser scan |
| IGM p881237 | *Eonatator* | *coellensis* | Paramo-Fonseca 2014 |
| PMU R 163 | *Eonatator* | *coellensis* | laser scan |
| OCP DEK-GE 112 | *Eremiasaurus* | *heterodontus* | LeBlanc et al. 2012 |
| UALVP 51744 | *Eremiasaurus* | *heterodontus* | LeBlanc et al. 2012 |
| MNHN 1891-14 | *Eremiasaurus* (?*Prognathodon*) | *mosasauroides* | structured-light scan |
| MHNM KHG 1231 | *Gavialimimus* | *almaghribensis* | Strong et al. 2020 |
| FHSM VP 13828 | *Globidens* | cf. *dakotensis* | laser scan |
| FMNH PR 846 | *Globidens* | *dakotensis* | structured-light scan |
| MHNM KHG 221 | *Globidens* | *simplex* | LeBlanc et al. 2019 |
| BYU 13082 | *Gnathomortis* | *stadtmani* | Lively et al. 2020 |
| IGF 14750-1 | *Goronyosaurus* | *nigeriensis* | Lingham-Soliar 1988 |
| EJ693 | *Haasiasaurus* | *gittelmani* | Polcyn et al. 1999 |
| EJ694 | *Haasiasaurus* | *gittelmani* | Polcyn et al. 1999 |
| UALVP 56123 | *Halisaurus* | *arambourgi* | Jimenez-Huidobro et al. 2017 |
| PMU R 163 | *Halisaurus* (*Eonatator*) | *sternbergii* | Bardet & Suberbiola 2001 |
| CDM 022 | *Kourisodon* | *puntledgensis* | Nicholls & Meckert 2002 |
| TMP 84-162-01 | *Latoplatecarpus* | *willistoni* | Konishi & Caldwell 2011 |
| NZGS CD 535 | *Moanasaurus* | *mangahouangae* | Bell et al. 1999 |
| OCP-DEK GE 303 | *Mosasaurus* | *beugei* | Bardet et al. 2004 |
| OCP-DEK GE 83 | *Mosasaurus* | *beugei* | Bardet et al. 2004 |
| MOR 006 | *Mosasaurus* | *conodon* | Ikejiri & Lucas 2014 |
| IRSNB R 12 | *Mosasaurus* | *hoffmannii* | structured-light scan; Lingham-Soliar 1995 |
| IRSNB R 3127 | *Mosasaurus* | *lemonnieri* | structured-light scan; Lingham-Soliar 2000 |
| KUVP 1034 | *Mosasaurus* | *missouriensis* | laser scan |
| TMP 2008.036.0001 | *Mosasaurus* | *missouriensis* | Konishi et al. 2014 |
| CM Zfr-1 | *Mosasaurus* | *mokoroa* | Welles & Gregg 1971 |
| MTM 2007 | *Pannoniasaurus* | *inexpectatus* | Makadi et al. 2012 |
| HMG 1528 | *Phosphorosaurus* | *ponpetelegans* | Konishi et al. 2015 |
| IRSNB R34 | *Phosphorosaurus* (*Halisaurus*) | *ortleibi* | Lingham-Soliar 1996 |
| FHSM VP 17017 | *Platecarpus* | *tympaniticus* | laser scan |
| FHSM VP 322 | *Platecarpus* | *tympaniticus* | photo |
| KGM 0035 | *Platecarpus* | *tympaniticus* | photo |
| KUVP 1007 | *Platecarpus* | *tympaniticus* | laser scan |
| KUVP 1031 | *Platecarpus* | *tympaniticus* | scan |
| KUVP 4862 | *Platecarpus* | *tympaniticus* | scan |
| LACM 128319 | *Platecarpus* | *tympaniticus* | Lindgren et al. 2010 |
| PMU | *Platecarpus* | *tympaniticus* | photo |
| FHSM VP 2116 | *Plesioplatecarpus* | *planifrons* | laser scan |
| FHSM VP 2181 | *Plesioplatecarpus* | *planifrons* | laser scan |
| FHSM VP 2296 | *Plesioplatecarpus* | *planifrons* | laser scan |
| UCMP 137249 | *Plesiotylosaurus* | *crassidens* | Lindgren 2009 |
| IRSNB 3101 | *Plioplatecarpus* | *houzeaui* | Lingham-Soliar 1994 |
| IRSNB 3130 | *Plioplatecarpus* | *houzeaui* | Lingham-Soliar 1994 |
| IRSNB R 35 | *Plioplatecarpus* | *houzeaui* | Lingham-Soliar 1994 |
| IRSNB R 37 | *Plioplatecarpus* | *houzeaui* | Lingham-Soliar 1994 |
| IRSNB R 38 | *Plioplatecarpus* | *marshi* | Lingham-Soliar 1994 |
| IRSNB R 39 | *Plioplatecarpus* | *marshi* | Lingham-Soliar 1994 |
| MOR 1062 | *Plioplatecarpus* | *peckensis* | Cuthbertson & Holmes 2015 |
| NMC 11835 | *Plioplatecarpus* | *primaevus* | Holmes 1996 |
| NMC 11840 | *Plioplatecarpus* | *primaevus* | Holmes 1996 |
| NMC 21854 | *Plioplatecarpus* | *primaevus* | Holmes 1996 |
| NMC P 1756.1 | *Plioplatecarpus* | *primaevus* | Holmes 1996 |
| TATE V 0087 | *Plioplatecarpus* | sp. | laser scan |
| UCMP 3718 | *Plotosaurus* | *bennisoni* | CT scan (Digimorph) |
| NHNM.KH 262 | *Pluridens* | *serpentis* | Longrich et al. 2021 |
| OCP DEK-GE 548 | *Pluridens* | *serpentis* | Longrich et al. 2021 |
| NHMUK R 14153 | *Pluridens* | *walkeri* | Longrich et al. 2021 |
| HUJ.OR 100 | *Prognathodon* | *currii* | Christiensen & Bonde 2002 |
| TMP 2002.400.0001 | *Prognathodon* | *overtoni* | Konishi et al. 2011 |
| TMP 2007.034.0001 | *Prognathodon* | *overtoni* | Konishi et al. 2011 |
| NHMM 1998141 | *Prognathodon* | *saturator* | Dortangs et al. 2002 |
| IRSNB R 33b | *Prognathodon* | *solvayi* | laser scan |
| NZGS CD 531 | *Rikisaurus* | *tehoensis* | Wiffen 1990 |
| MPPS 42224 | *Romeosaurus* | *fumanensis* | Palci et al. 2013 |
| SMU 73056 | *Russellosaurus* | *coheni* | 2D+ scan; Polcyn & Bell 2005 |
| FHSM VP 13910 | *Selmasaurus* | *johnsoni* | laser scan |
| IAA 2000-JR-FSM-1 | *Taniwhasaurus* | *antarcticus* | Novas et al. 2002 |
| KHM N99-1014 | *Taniwhasaurus* | *oweni* | Caldwell et al. 2005 |
| NMNZ R 1532 | *Taniwhasaurus* | *oweni* | Caldwell et al. 2005 |
| NMNZ R 1536 | *Taniwhasaurus* | *oweni* | Caldwell et al. 2005 |
| MNHN 1999-9 | *Tethysaurus* | *nopscai* | Bardet et al. 2003 |
| FHSM VP 2209 | *Tylosaurus* | *kansasensis* | laser scan |
| FHSM VP 2295 | *Tylosaurus* | *nepaeolicus* | laser scan |
| MM V 95 | *Tylosaurus* | *pembinensis* | Bulland & Caldwell 2010 |
| AMNH FARB 221 | *Tylosaurus* | *proriger* | photo |
| FFHM 1997-10 | *Tylosaurus* | *proriger* | laser scan |
| FHSM VP 3 | *Tylosaurus* | *proriger* | laser scan |
| FMHN GEO 79878 | *Tylosaurus* | *proriger* | structured-light scan |
| GPIT RE 9422 | *Tylosaurus* | *proriger* | scan |
| KUVP 1032 | *Tylosaurus* | *proriger* | laser scan |
| KUVP 28705 | *Tylosaurus* | *proriger* | laser scan |
| RSM P 2588.1 | *Tylosaurus* | *sasckatchwanensis* | Jimenez-Huidoboro et al. 2018 |
| IRSNB R23 | *Tylosaurus* (*Hainosaurus*) | *bernardi* | laser scan |
| BRV-68 | *Yaguarasaurus* | *columbianus* | Paramo-Fonseca 2000 |

2. **Functional Ratios**

| **Trait** | **Function** | **Measurement** |
| --- | --- | --- |
| ***Feeding*** | | |
| Depressor Lever Arm Ratio | Proxy for mechanical advantage of the *depressor mandibularis* | Retroarticular process length ÷ Mandible Length (articulation to tip) |
| Adductor Lever Arm Ratio | Proxy for mechanical advantage of the *adductor externus* | Articulation to apex of coronoid ÷ Mandible Length (articulation to tip) |
| Functional Toothrow | Describes proportion of the jaw available for prey capture | Dentigerous mandible length ÷ Mandible Length (articulation to tip) |
| Functional Jaw Robusticity | Robusticity of the functional jaw; proxy for jaw strength | Jaw Depth beneath centre of toothrow ÷ Mandible Length (articulation to tip) |
| Supratemporal Fenestra Area | Proxy for cross-sectional area of combined adductor musculature | (STF length x STF width) ÷ 2  Mandible Length (articulation to tip) ^2^ |
| Longirostry | Describes elongation of the pre-orbital snout (hydrodynamic potential of the snout) | Pre-orbital snout length ÷ Mandible Length (articulation to tip) |
| Gullet | Proxy for the width of the gullet (largest potential prey; volume of water to be expelled from mouth) | Pre-orbital snout width ÷ Mandible Length (articulation to tip) |
| Tooth Crown Shape | Proxy for tooth narrowing; informs on potential food items (hard vs. soft) | Mean tooth height ÷ mean tooth anteroposterior length |
| Tooth Blade Shape | Describes dental compression; conical vs. blade-like teeth | Mean tooth anteroposterior length ÷ mean tooth labiolingual width |
| Crown Curvature | Describes dental curvature, proxy for potential prey items | Mean crown tip offset ÷ mean tooth height |
| ***Sensory Perception*** | | |
| Relative Nares Size | Proxy for volume of air possible to inhale in single breath | Length of narial opening ÷ Mandible Length (articulation to tip) |
| Nares Retraction | Proxy describing the ease of which a breath can be taken during steady state swimming | Distance from centroid of narial opening to anterior snout ÷ Mandible Length (articulation to tip) |
| Orbit Size | Proxy for eye size (and therefore potential visual acuity) | Mean diameter of orbit x π .  Mandible Length (articulation to tip) |
| Pupil Size | Defined as the space available between the sclerotic rings which allows light to enter the pupil | (Sclerotic opening W + Sclerotic opening L)  2 |
| Tympanic Resonator | Proxy for the area of the quadrate available as a sound resonator | (Tympanic conch H x Tympanic conch W) ÷ 2  Mandible Length (articulation to tip) ^2^ |
| Premaxilla Elongation | Defines the shape of the premaxilla; proxy for anterior hydrodynamics and area available for pressure sensation | Length of premaxilla (from articulation with maxilla) ÷ maximum width of premaxilla |
| Parietal Foramen | Defines the length of the parietal foramen; proxy for relative size of the pineal eye | Length of parietal foramen ÷ Mandible Length (articulation to tip) |

3. **Species Coverage**

List of species used in this study. Number of specimens (No. spec.) per species is listed alongside the percentage of traits which were scored for each species for ‘feeding’ (% feeding) dataset and entire bauplan (% bauplan; cranial + postcranial traits) datasets. * based on reconstruction / part-reconstruction in literature; Y = 3D scanned

| **Genus** | **Species** | **No. spec.** | **3D** | **% feeding** |
| --- | --- | --- | --- | --- |
| *Angolasaurus* | *bocagei* | 1 |  | 90 |
| *Carinodens* | *belgicus* | 1 |  | 30 |
| *Clidastes* | *propython* | 9 | Y | 100 |
| *Ectenosaurus* | *clidastoides* | 1 | Y | 100 |
| *Eonatator* | *coellensis* | 1 |  | 80 |
| *Eonatator* (*Halisaurus*) | *sternbergii* | 1 | Y | 70 |
| *Eremiasaurus* | *heterodontus* | 2 |  | 90 |
| *Eremiasaurus* | *mosasauroides* | 1 | Y | 40 |
| *Gavialimimus* | *almaghribensis* | 1 |  | 100 |
| *Globidens* | *dakotensis* | 2 | Y | 70 |
| *Globidens* | *simplex* | 1 |  | 70 |
| *Gnathomortis* | *stadtmani* | 1 |  | 60 |
| *Goronyosaurus* | *nigeriensis* | 1 |  | 90* |
| *Haasiasaurus* | *gittelmani* | 1 |  | 60 |
| *Halisaurus* | *arambourgi* | 1 |  | 90 |
| *Kourisodon* | *puntledgensis* | 1 |  | 60 |
| *Latoplatecarpus* | *willistoni* | 1 |  | 100 |
| *Moanasaurus* | *mangahouanganae* | 1 |  | 100 |
| *Mosasaurus* | *conodon* | 2 |  | 90 |
| *Mosasaurus* | *hoffmannii* | 1 | Y | 100 |
| *Mosasaurus* | *beaugei* | 2 |  | 70 |
| *Mosasaurus* | *lemonnieri* | 2 | Y | 100 |
| *Mosasaurus* | *missouriensis* | 2 | Y | 100 |
| *Mosasaurus* | *mokoroa* | 1 |  | 60 |
| *Pannoniasaurus* | *inexpectatus* | 1 |  | 80 |
| *Phosphorosaurus* | *ortleibi* | 1 |  | 70* |
| *Phosphorosaurus* | *ponpetelegans* | 1 |  | 80 |
| *Platecarpus* | *coryphaeus* | 3 | Y | 100 |
| *Platecarpus* | *tympaniticus* | 5 | Y | 100 |
| *Plesioplatecarpus* | *planifrons* | 3 | Y | 100 |
| *Plesiotylosaurus* | *crassidens* | 1 |  | 100 |
| *Plioplatecarpus* | *houzeaui* | 2 |  | 90 |
| *Plioplatecarpus* | *marshi* | 1 |  | 80 |
| *Plioplatecarpus* | *peckensis* | 1 |  | 60 |
| *Plioplatecarpus* | *primaevus* | 4 |  | 100 |
| *Plioplatecarpus* | *sp.* | 1 | Y | 60 |
| *Plotosaurus* | *bennisonni* | 2 | Y | 100 |
| *Pluridens* | *walkeri* | 1 |  | 40 |
| *Prognathodon* | *currii* | 1 |  | 100 |
| *Prognathodon* | *overtoni* | 2 |  | 90 |
| *Prognathodon* | *saturator* | 1 |  | 60 |
| *Prognathodon* | *solvayi* | 1 | Y | 100 |
| *Rikisaurus* | *tehoensis* | 1 |  | 100 |
| *Romeosaurus* | *fumanensis* | 1 |  | 70 |
| *Russellosaurus* | *coheni* | 1 | Y | 100 |
| *Selmasaurus* | *johnsoni* | 1 | Y | 100 |
| *Taniwhasaurus* | *antarcticus* | 1 |  | 80 |
| *Taniwhasaurus* | *oweni* | 1 |  | 60 |
| *Tethysaurus* | *nopcsai* | 1 | Y | 90 |
| *Tylosaurus* | *bernardi* | 1 | Y | 100 |
| *Tylosaurus* | *kansasensis* | 1 | Y | 90 |
| *Tylosaurus* | *dyspelor* | 2 |  | 100 |
| *Tylosaurus* | *nepaeolicus* | 1 | Y | 100 |
| *Tylosaurus* | *pembinensis* | 1 |  | 100 |
| *Tylosaurus* | *proriger* | 5 | Y | 100 |
| *Tylosaurus* | *sasckatchwanensis* | 1 |  | 90 |
| *Yaguarasaurus* | *colombianus* | 1 |  | 70 |

4. **Supplementary Figures**

**Supplementary Figure S1:** Functional ecomorphospace (based on PCoA axes) and size distribution in mosasaurids.

**Supplementary Figure S2**: Phylogenetic clade disparity through Maastrichtian

5. **Supplementary Tables**

**Supplementary Table S1:** Comparisons of significant differences (Wilcoxon test) between Maastrichtian time bins under different bootstrapping regimes

**Supplementary Table S2:** Significant differences (Wilcoxon test) in clade disparity between Maastrichtian time bins

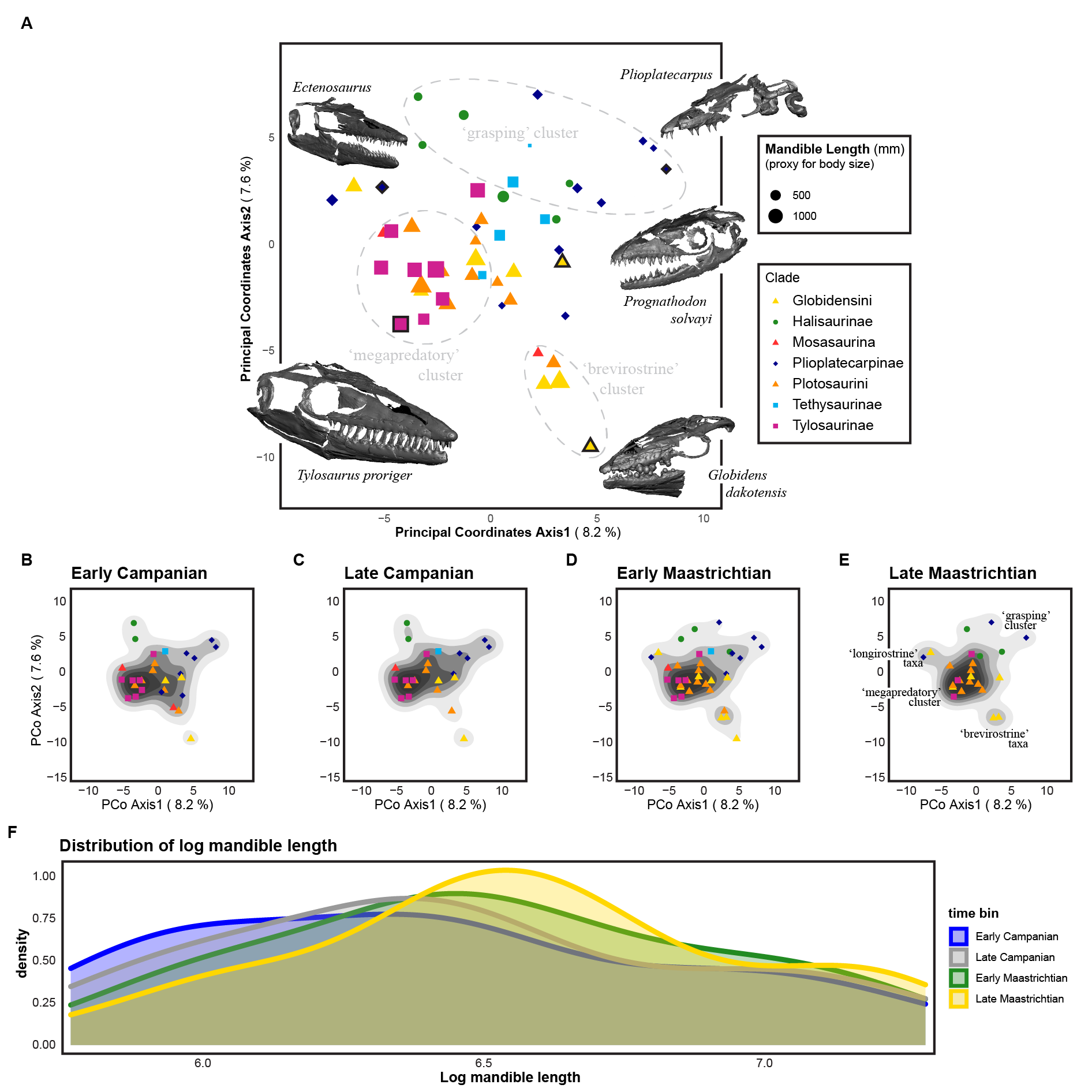

**Supplementary Figure S1.** Functional ecomorphospace and size distribution in mosasaurids. Functional ecomorphospace occupation (based on PCoA axes 1 and 2) by all mosasaurids in the sample (**A**) with ecomorphological clusters and representative 3D models of skulls. The first two PCo Axes account for only 15.8% of ecomorphological variation in the sample; by contrast, the first two NMDS axes (Figure 2A-E, main text) account for all ecomorphological variation in a non-metric framework. Data points outlined in bold represent the placement of exemplar skulls; data point size represents relative skull size (based on mandible length). Functional ecomorphospace for each time bin through the Campanian-Maastrichtian (**B-E**) demonstrating changes in density and isolation of ecomorphological clusters in Late Maastrichtian (**E**). Size distribution of mosasaurids through the Campanian-Maastrichtian (**F**) demonstrating shifts in the density of small and mid-sized mosasaurids from Early Campanian (ECam) to Late Maastricthian (LMaa) (after Figure 2, main text).

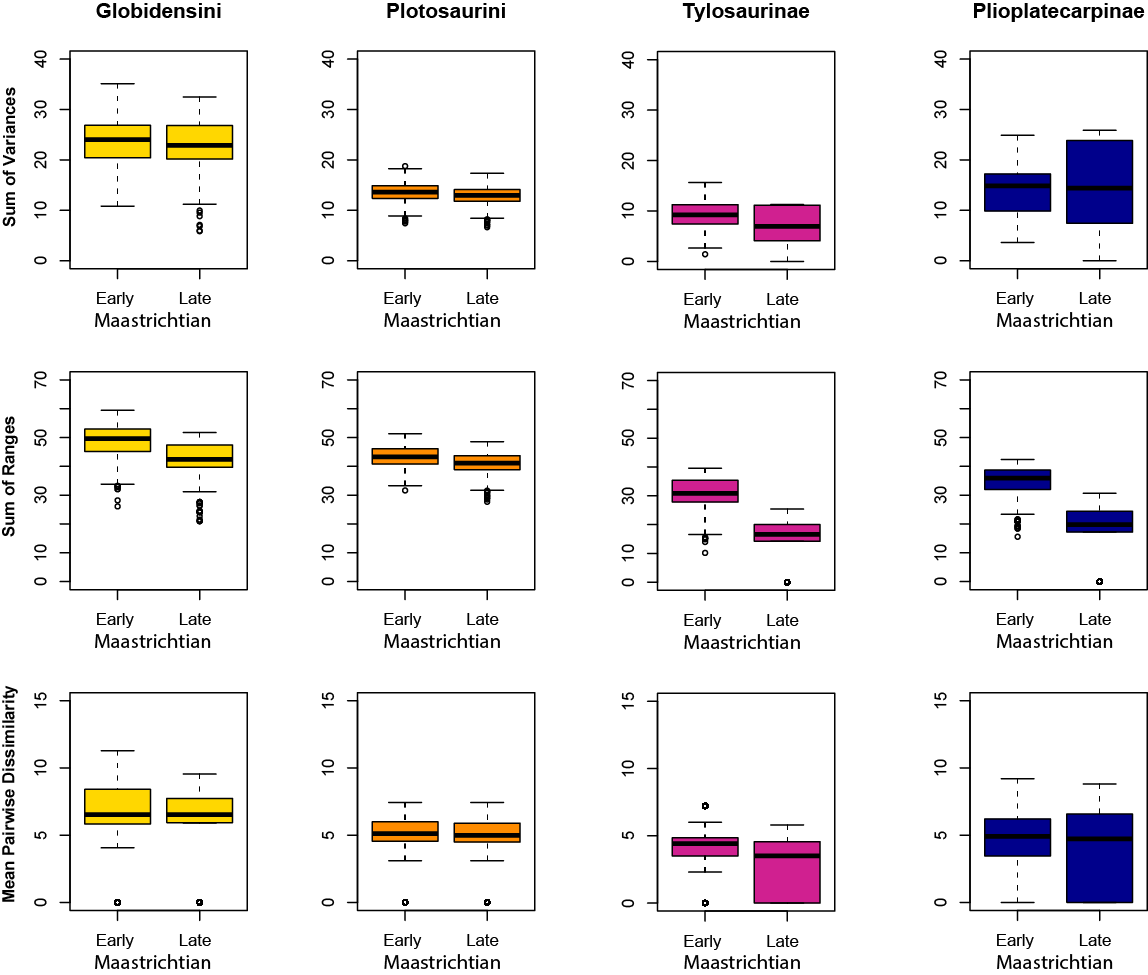

**Supplementary Figure S2.** Disparity pattern through the Maastrichtian time bins at the phylogenetic clade level. Three different disparity metrics were used: sum of variances (SoV), sum of ranges (SoR) and mean pairwise dissimilarity (MPD), with Bonferroni corrections for multiple comparisons. Halisaurines, tethysaurines and indeterminate mosasaurines were excluded due to low taxon counts in the sample for the Maatrichtian. Significance values charted in Supplementary Table S2.

**Supplementary Table S1**. Comparisons of significant differences (Wilcoxon test) between Early- and Late Maastrichtian time bins under different bootstrapping regimes (1000bs, 500bs, 250bs, 100bs), used as a sensitivity test for the effect of bootstrapping on our results. Sum of variances disparity metric used, with a Bonferroni correction for multiple comparisons. Provincial regions: WIS = Western Interior Seaway, NTP = Norther Tethys Province, STP = Southern Tethys Province, WP = Weddellian Province. Wilcoxon test statistic (W) reported alongside p-values (alpha set at ≤0.01; significant values shaded grey).

| **Region** | **1000 bs** | | **500 bs** | | **250 bs** | | **100 bs** | |
| --- | --- | --- | --- | --- | --- | --- | --- | --- |
|  | **W** | **p** | **W** | **p** | **W** | **p** | **W** | **p** |
| Global | 627165 | <0.001 | 167809 | <0.001 | 40800 | <0.001 | 6483 | 0.002 |
| WIS | 955795 | <0.001 | 239684 | <0.001 | 59010 | <0.001 | 9799 | <0.001 |
| NTP | 915984 | <0.001 | 229700 | <0.001 | 57593 | <0.001 | 9051 | <0.001 |
| STP | 567301 | <0.001 | 142775 | <0.001 | 35934 | <0.001 | 5489 | 0.232 |
| WP | 718336 | <0.001 | 178093 | <0.001 | 45499 | <0.001 | 6746 | <0.001 |

**Supplementary Table S2**. Significant differences (Wilcoxon test) between phylogenetic clade disparity from Early- and Late Maastrichtian (1000bs) under different disparity metrics. Sum of variances (SoV), sum of ranges (SoR) and mean pairwise dissimilarity (MPD) disparity metrics were used, with false-discovery rate corrections for multiple comparisons. Halisaurines, tethysaurines and indeterminate mosasaurines were excluded due to low taxon counts in the sample for the Maatrichtian. Wilcoxon test statistic (W) reported alongside respective p-values (alpha set at ≤0.01; significant values shaded grey).

| **Phylogenetic Clade** | **Sum Variance** | | **Sum Ranges** | | **Mean PD** | |
| --- | --- | --- | --- | --- | --- | --- |
|  | **W** | **p** | **W** | **p** | **W** | **p** |
| Globidensini (Mosasaurina) | 531619 | 0.01 | 816983 | <0.01 | 205692474 | <0.01 |
| Plotosaurini (Mosasaurina) | 600857 | <0.01 | 669617 | <0.01 | 859607915 | <0.01 |
| Tylosaurinae (Russellosaurina) | 606172 | <0.01 | 954784 | <0.01 | 26319339 | <0.01 |
| Plioplatecarpinae (Russellosaurina) | 425997 | <0.01 | 943718 | <0.01 | 31472304 | 0.94 |

**Supplementary Table S3**. Significant differences (Wilcoxon test) in γ-disparity between sequential time bins from Early Campanian to Late Maastrichtian (1000bs) under different disparity metrics. Sum of variances (SoV), sum of ranges (SoR) and mean pairwise dissimilarity (MPD) disparity metrics were used, with Bonferroni rate corrections for multiple comparisons. Wilcoxon test statistic (W) reported alongside respective p-values (alpha set at ≤0.01; significant values shaded grey). ECam = Early Campanian; LCam = Late Campanian

| **Disparity Metric** | **ECam-LCam** | | **LCam-EMaa** | | **EMaa-LMaa** | |
| --- | --- | --- | --- | --- | --- | --- |
|  | **W** | **p** | **W** | **p** | **W** | **p** |
| SoV | 485369 | 1.00 | 267611 | <0.01 | 637287 | <0.01 |
| SoR | 774941 | <0.01 | 4707 | <0.01 | 970578 | <0.01 |
| MPD | 51594741224 | <0.01 | 76877712744 | 1.00 | 89672062471 | <0.01 |
